# Mapping ribonucleotides embedded in genomic DNA to single-nucleotide resolution using Ribose-Map

**DOI:** 10.1101/2020.08.27.267153

**Authors:** Alli L. Gombolay, Francesca Storici

**Affiliations:** School of Biological Sciences, Georgia Institute of Technology, 950 Atlantic Drive NW, Atlanta, GA 30332-0230 USA

## Abstract

Ribose-Map is a user-friendly, standardized bioinformatics toolkit for the comprehensive analysis of ribonucleotide sequencing experiments. It allows researchers to map the locations of ribonucleotides in DNA to single-nucleotide resolution and identify biological signatures of ribonucleotide incorporation. In addition, it can be applied to data generated using any currently available high-throughput ribonucleotide sequencing technique, thus standardizing the analysis of ribonucleotide sequencing experiments and allowing direct comparisons of results. This protocol describes in detail how to use Ribose-Map to analyze raw ribonucleotide sequencing data, including preparing the reads for analysis, locating the genomic coordinates of ribonucleotides, exploring the genome-wide distribution of ribonucleotides, determining the nucleotide sequence context of ribonucleotides, and identifying hotspots of ribonucleotide incorporation. Ribose-Map does not require background knowledge of ribonucleotide sequencing analysis and assumes only basic command-line skills. The protocol requires less than 3 hr of computing time for most datasets and about 30 min of hands-on time.

## INTRODUCTION

High-throughput sequencing techniques have been developed recently to capture the locations of the most abundant type of non-canonical nucleotides in the genome, ribonucleoside monophosphates (rNMPs). These sequencing techniques include ribose-seq ^1^, emRiboSeq ^2^, RHII-HydEn-seq ^3^, Alk-HydEn-seq ^4^, and Pu-seq ^5^. Each of these rNMP sequencing techniques can capture potentially millions of rNMPs, generating large, complex datasets requiring software that can accurately and efficiently transform raw sequencing data into biologically meaningful results. In addition, such software should be able to accommodate data generated from any currently available high-throughput rNMP sequencing technique to standardize the analysis of rNMP sequencing experiments. Such standardization would facilitate direct comparisons of results obtained using different rNMP sequencing techniques to assess their reproducibility. The analysis pipeline for rNMP sequencing data can be divided into three main tasks: (i) alignment of sequencing reads to the reference genome, (ii) mapping the genomic coordinates of rNMPs, and (iii) identifying biological signatures of rNMP incorporation. To accurately and efficiently accomplish these tasks for any type of rNMP sequencing data, we developed the Ribose-Map bioinformatics toolkit ^6^, which includes the Alignment, Coordinate, Sequence, Distribution, and now also the Composition and Hotspot Modules. In particular, the Alignment Module aligns the data to the reference genome, and then the Coordinate Module uses the alignment results to map the genomic coordinates of rNMPs for the corresponding rNMP sequencing technique. Collectively, the Composition, Sequence, Distribution, and Hotspot Modules identify biological signatures of rNMP incorporation. The Sequence Module determines the nucleotide sequence context of rNMPs, while the Distribution Module assesses the genome-wide distribution of rNMPs. The newly added Composition Module determines the rNMP composition (i.e., frequencies of r[A, C, G, U]MP normalized to those of the reference genome), and the Hotspot Module identifies genomic sites that are enriched for rNMP incorporation and their consensus sequence.

Ribose-Map is a proven technique that we previously applied to characterize the biological signatures of rNMP incorporation in rNMP sequencing libraries of different strains, genotypes, and species of yeast prepared using two markedly different rNMP sequencing techniques, ribose-seq and emRiboSeq ^7^. Here, we outline a detailed protocol for Ribose-Map and demonstrate how to use each of its modules to analyze yeast rNMP sequencing data generated using ribose-seq and emRiboSeq. The steps outlined in this protocol could be easily adapted to analyze rNMP sequencing data derived from any organism with a sequenced reference genome and generated using any rNMP sequencing technique. Ribose-Map is fully documented and actively maintained by the developers at https://github.com/agombolay/ribose-map. By following this protocol, researchers can gain insight into the biological mechanisms that regulate the presence of rNMPs in DNA and their effects on genome stability, DNA metabolism, and disease.

### Applications of Ribose-Map

Researchers can use Ribose-Map to study the biological signatures of rNMP incorporation in any genome of interest provided a reference genome is available. To facilitate comparisons among rNMP sequencing data, not only those derived from different rNMP libraries prepared using the same rNMP sequencing technique, but particularly those derived from libraries prepared using different rNMP sequencing techniques, Ribose-Map is standardized to allow researchers to directly compare the results of any rNMP sequencing experiment regardless of the rNMP sequencing technique used. In addition to rNMPs, Ribose-Map can also be used to study the biological signatures of any single-nucleotide genomic coordinates of interest, such as singlenucleotide polymorphisms. To perform such an analysis, the user should bypass the Alignment and Coordinate Modules of Ribose-Map and directly input the coordinates provided in BED file format into the Composition, Sequence, Distribution, and/or Hotspot Modules.

### Comparison of Ribose-Map with Current Methods

Ribose-Map is currently the only available rNMP mapping computational toolkit that can process and analyze rNMP sequencing data generated using any rNMP sequencing technique. In contrast to Ribose-Map, the computational methods applied in previously published rNMP sequencing studies are customized to analyze rNMP sequencing data generated using only the rNMP sequencing technique applied in that particular study, and thus are not standardized. In addition, these current computational methods do not output the genomic coordinates of rNMP sites, depend on proprietary software, do not produce visualizations of the data, and/or are limited in scope. In contrast, Ribose-Map processes and analyzes rNMP sequencing data for any rNMP sequencing technique, outputs the genomic coordinates of rNMPs, depends upon only opensource software, and produces summary datasets and visualizations for the nucleotide composition, DNA sequence context, genome-wide distribution, and hotspots of rNMPs. Ribose-Map also generates well-formatted output files (e.g., BED/BedGraph) containing the genomic coordinates of rNMPs that can be directly input into other commonly used bioinformatics software, such as JBrowse ^8^, UCSC genome browser ^9^, and Bedtools ^10^.

### Experimental Design

The Ribose-Map bioinformatics toolkit consists of six analytical modules that collectively align rNMP sequencing data to the reference genome, calculate the single-nucleotide genomic coordinates of rNMPs, and uncover biological signatures of rNMP incorporation in the genome of interest. **Figure 1** shows the input/output of the six modules of Ribose-Map.

**Figure 1.**
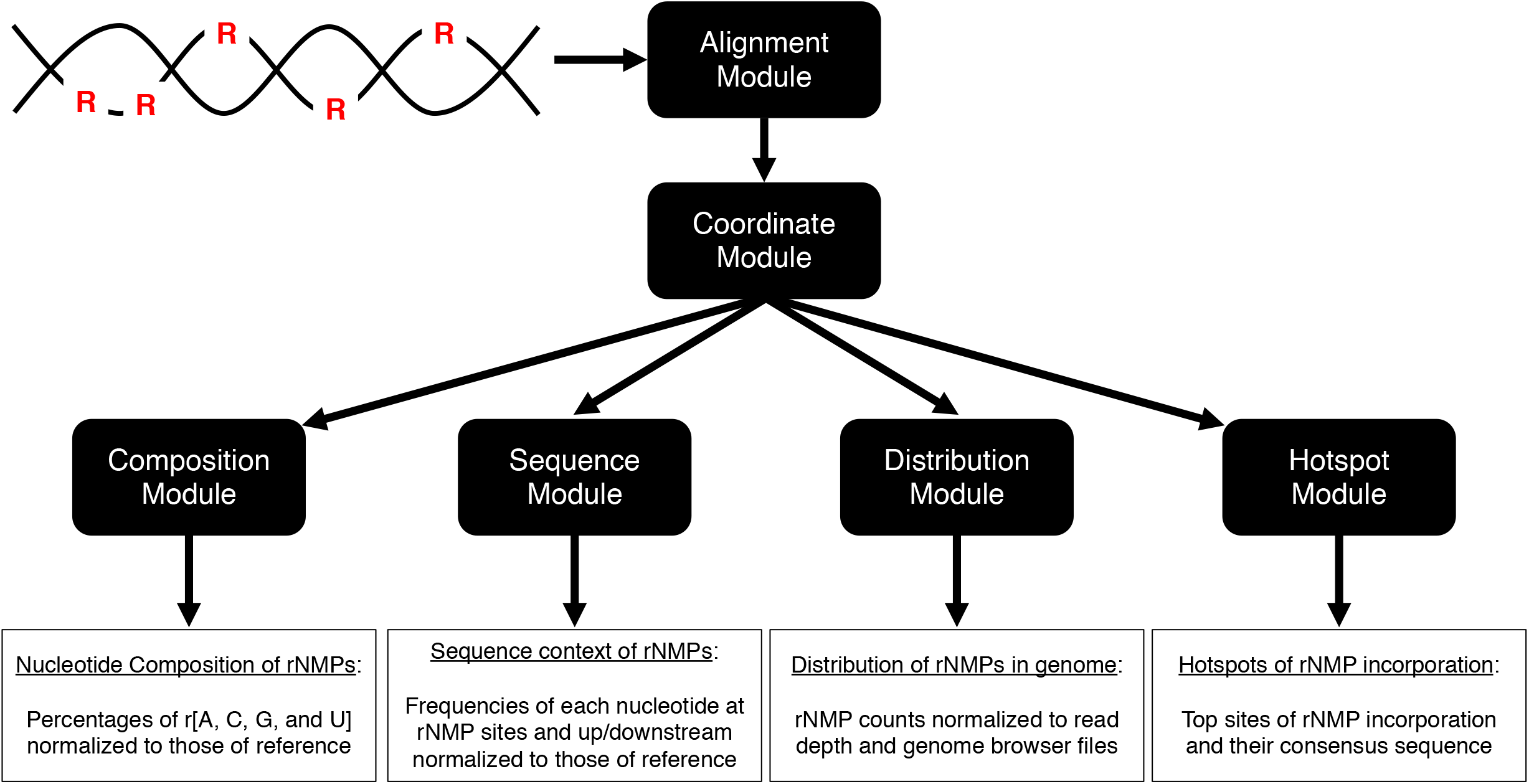
Input/output of Ribose-Map’s modules.

### Read Alignment with Alignment Module

Analysis of rNMP sequencing data begins by aligning reads to the corresponding reference genome to locate their genomic positions. Ribose-Map’s Alignment Module aligns single- or paired-end sequencing reads to the reference genome using Bowtie 2. The Alignment Module also processes reads for unique molecular identifiers (UMIs) and molecular barcodes using UMI-tools ^11^ and seqtk (https://github.com/lh3/seqtk), respectively. If PCR was used during library preparation, a UMI is critical to remove PCR duplicates from the data. In addition, a molecular barcode is helpful to separate real data from artifacts. If necessary, the reads should be processed with Cutadapt ^12^ or similar data cleaning software to remove low-quality base calls and sequencing adapters prior to alignment.

### Locating rNMPs with the Coordinate Module

During rNMP sequencing library preparation, rNMPs are tagged relative to the 5’ end of the sequencing read (read 1 if paired-end). **Figure 2** compares the rNMP position relative to the 5’ end of the sequencing read for ribose-seq, emRiboSeq, RHII-HydEn-seq, Alk-HydEn-seq, and Pu-seq data. To calculate the genomic coordinates of rNMPs, the Coordinate Module converts the BAM file containing the aligned positions of sequencing reads output by the Alignment Module into a BED file containing the zero-based start/end positions of the aligned reads using BEDTools. Then, the Coordinate Module uses customized shell commands to calculate the single-nucleotide genomic coordinates of rNMPs relative to the 5’ positions of the sequencing read for the corresponding rNMP sequencing technique. **Table 1** shows the genomic arithmetic used to calculate the genomic coordinates of rNMPs for ribose-seq, emRiboSeq, RHII-HydEn-seq, Alk-HydEn-seq, and Pu-seq data. Although unlikely, contamination or inefficiencies during library preparation could result in sequencing reads that align to the 5’-most ends of chromosomes, resulting in rNMP coordinates that are located beyond the chromosome ends and thus biologically meaningless. Therefore, once the genomic coordinates of rNMPs have been calculated, the Coordinate Module performs quality control on the rNMP coordinates prior to saving them to the output BED file to prevent computational issues during downstream analyses.

**Figure 2.**
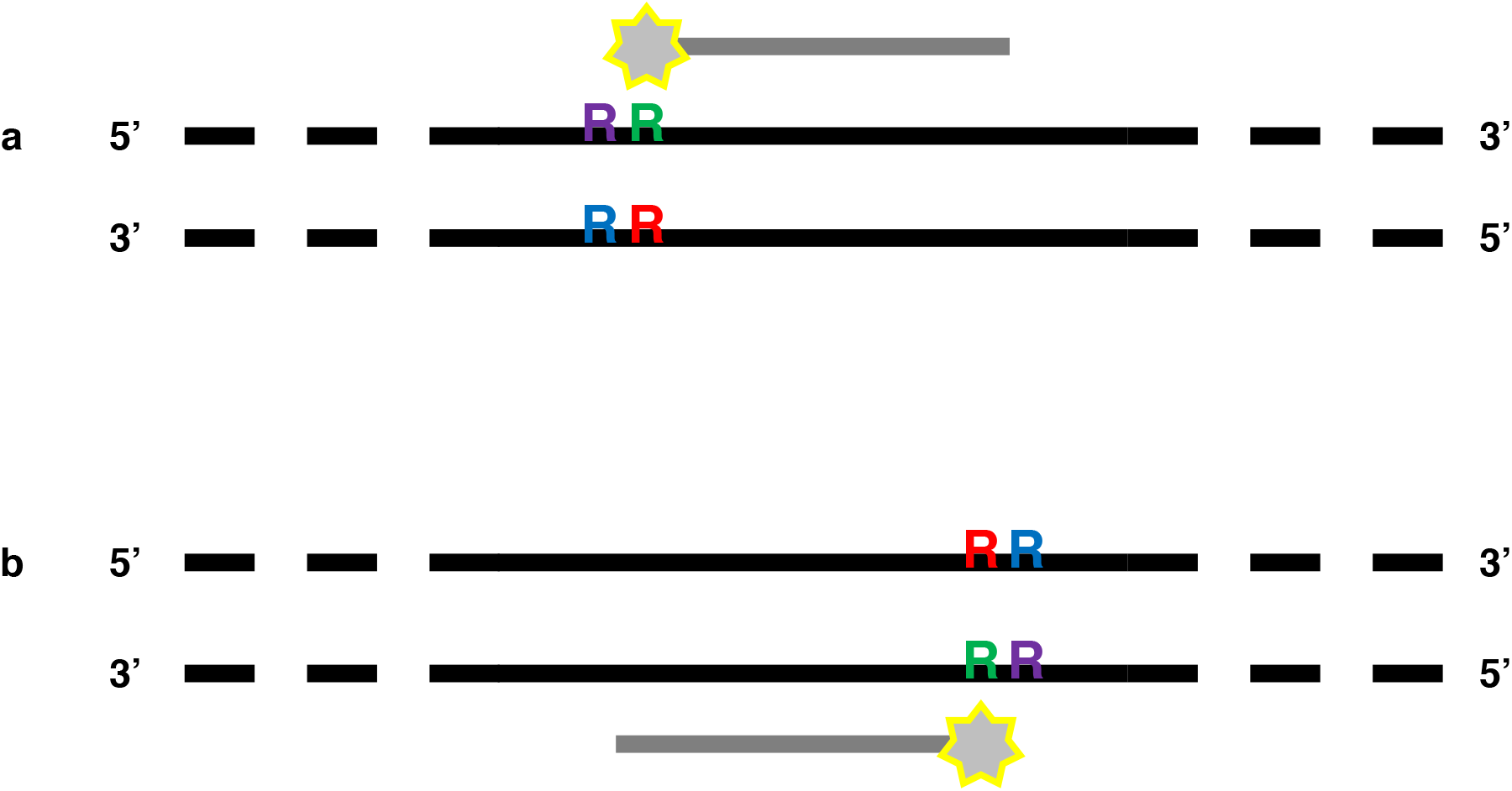
Comparison of positions of an rNMP (‘R’) embedded in genomic DNA (black lines) relative to the tagged 5’-most nucleotide (star) of a sequencing read (grey line) for reads that align to the (a) Watson strand and (b) Crick strand for ribose-seq (red ‘R’), emRiboSeq (blue ‘R’), RHII-HydEn-seq (green ‘R’), Alk-HydEn-seq (purple ‘R’), and Pu-seq (purple ‘R’). For ribose-seq, the rNMP (red ‘R’) is the reverse complement of the tagged nucleotide; for emRiboSeq, the rNMP (blue ‘R’) is one nucleotide downstream from the reverse complement of the tagged nucleotide; for RHII-HydEn-seq, the rNMP (green ‘R’) is in the same position as the tagged nucleotide; for Alk-HydEn-seq and Pu-seq, the rNMP (purple ‘R’) is one nucleotide upstream from the tagged nucleotide.

**Table 1.** Genomic arithmetic used to calculate the genomic coordinates of rNMPs for ribose-seq, emRiboSeq, RHII-HydEn-seq, Alk-HydEn-seq, and Pu-seq.

|  | Reads aligned to + DNA Strand |  |  | Reads aligned to – DNA strand |  |  |
| --- | --- | --- | --- | --- | --- | --- |
|  | Start | End | Strand | Start | End | Strand |
| **ribose-seq** | S | S + 1 | – | E – 1 | E | + |
| **emRiboSeq** | S – 1 | S | – | E | E + 1 | + |
| **RHII-HydEn-seq** | S | S + 1 | + | E – 1 | E | – |
| **Alk-HydEn-seq**<br>**and Pu-seq** | S – 1 | S | + | E | E + 1 | – |

Start represents the 0-based start coordinate of rNMP; End represents the 0-based end coordinate of rNMP; S represents the 0-based start coordinate of a read; E represents the 0-based end coordinate of a read; Strand represents the Strand of rNMP; + represents the Watson strand and – represents the Crick strand.

### Studying biological signatures of rNMP incorporation using the Composition, Sequence, Distribution, and Hotspot Modules

Once the genomic coordinates of rNMPs have been calculated, researchers can input these coordinates into Ribose-Map’s Composition, Sequence, Distribution, and Hotspot Modules to uncover biological signatures of rNMP incorporation in the genome of interest. In particular, researchers can use these modules to explore four important questions: (i) What is the genomewide distribution of rNMP incorporation? (ii) Are there biases in the type of rNMP nucleotide incorporated into genomic DNA? (iii) Is rNMP incorporation influenced by the surrounding DNA sequence context? (iv) Are there hotspots of rNMP incorporation?

The Sequence Module calculates and plots the frequencies of the nucleotides at the sites of rNMP incorporation and up/downstream from those sites. These frequencies are normalized to those of the reference genome. The Distribution Module calculates and plots the per-nucleotide coverage of rNMP at each position in the genome and normalizes the coverage to the total read count to allow for direct comparison among libraries of different sequencing depth. The Distribution Module also creates BedGraph files that can be directly uploaded to a genome browser, such as the University of California, Santa Cruz Genome Browser (http://genome.ucsc.edu/), as custom annotation tracks. In addition to the Sequence and Distribution Modules, we have now added two additional modules to the Ribose-Map toolkit, the Composition Module and the Hotspot Module. The Composition Module calculates the percentages of r[A, C, G, U]MP and normalizes the percentages to those of the reference genome. The Hotspot Module identifies the top 1% (or other %, as desired) most abundant sites of rNMP incorporation and uses MEME to identify any consensus sequences of those sites.

To demonstrate the potential of Ribose-Map, we consider previously published rNMP sequencing data derived from ribonuclease (RNase) H2-defective strains of *Saccharomyces cerevisiae* using ribose-seq library SRR11364933 (strain background E134) ^7^ and emRiboSeq library SRR1734967 (strain background (Δl−2)l-7BYUNI300) ^2^.

### Expertise Needed to Implement the Protocol

This protocol assumes experience with the Unix/Linux command-line interface. Users should be able to run programs from the command-line and edit text files in the Unix/Linux environment.

### Limitations of the Protocol

Since Ribose-Map’s Alignment Module employs the short-read aligner, Bowtie 2, longer reads generated using third-generation sequencing technology, such as Pacific Biosciences, should not be input into the Alignment Module. However, after aligning longer reads using an appropriate software, the user can directly input the aligned reads into Ribose-Map’s Coordinate Module.

## MATERIALS

### EQUIPMENT

- Ribose-Map GitHub repository (https://github.com/agombolay/ribose-map)
- Data (rNMP sequencing reads, reference genome files, and configuration file)
- Anaconda3 (https://docs.conda.io/en/latest/)
- Git version-control system (https://git-scm.com/)
- Hardware (64-bit computer running Linux or Mac OS X; 4 GB of RAM)

#### Equipment Set-up

#### Installing Software

Ribose-Map utilizes a combination of open-source bioinformatics software and custom Linux and R commands. To facilitate proper software versioning and reduce set-up time, we recommend using Anaconda3 (see EQUIPMENT) to create a software environment in which to install the required software. The YAML file required to create such an environment for Linux or MacOSX is provided in the ribose-map/lib directory of the Ribose-Map GitHub repository. Instructions on installing the git version-control system are provided at https://git-scm.com/. All commands shown below assume a Bash Shell and should be run via the Unix/Linux command-line (indicated by “$’).

Clone Ribose-Map repository from GitHub:

$ git clone https://github.com/agombolay/ribose-map.git

Create conda software environment:

$ conda env create --name ribosemap_env --file ribose-map/lib/ribosemap.yaml

#### Required Data

Ribose-Map requires (i) a FASTQ file of single- or paired-end rNMP sequencing data (ribose-seq, emRiboSeq, RHII-HydEn-seq, Alk-HydEn-seq, or Pu-seq), (ii) a FASTA file of the reference genome nucleotide sequence, (iii) a file containing the reference genome chromosome sizes (extension .chrom.sizes), (iv) Bowtie 2 indexes for the reference genome (extension .bt2), and (v) a configuration (config) file. The FASTQ file used in this protocol can be downloaded from NCBI’s Short Read Archive via BioProject PRJNA613920. The FASTA and .chrom.sizes files for many genomes, including the 2008 build of the *S. cerevisiae* genome (sacCer2) used in this protocol, are available at the UCSC genome browser website (https://genome.ucsc.edu/). Instructions on building the Bowtie 2 indexes from the FASTA file are provided below. The config files used in this protocol are provided in the ribose-map/config directory of the Ribose-Map GitHub repository and can be adapted for other rNMP sequencing experiments as needed. **Table 2** lists the parameters that should be specified in the configuration file.

**Table 2.** Parameters for configuration file.

| Parameters | Examples |
| --- | --- |
| **rNMP Sequencing Data** |  |
| Sample name for labelling output | sample='SRR123' |
| Read 1 of rNMP sequencing data | read1='/local/path/SRR123_1.fastq' |
| Read 2 of rNMP sequencing data | read2='/local/path/SRR123_2.fastq' |
| rNMP sequencing technique | technique='ribose-seq' |
| **UMI and Molecular Barcode** |  |
| Molecular barcode in read 1 | barcode='ATC' |
| UMI/barcode pattern in read 1 | pattern='NNNXXXNNN' |
| **Reference Genome** |  |
| Fasta file of reference genome | fasta='/local/path/genome.fa' |
| Basename of Bowtie 2 indexes | basename='/local/path/genome' |
| Genomic units to analyze individually | other='chrM chrX chrY' |
| **Software Information** |  |
| Local Ribose-Map repository | repository='path/ribose-map' |
| Percentile for Hotspot Module | percentile='0.99' |
| Number of computing threads | threads='4' |
| Minimum quality of alignment | quality='30' |

All parameters are required except 'barcode', 'pattern', 'read2', and 'other'. The number of 'N' characters in 'pattern' represents the length of the UMI sequence. The number of 'X' characters represents the length of the barcode sequence, and the position of these characters represents the position of the barcode relative to the UMI. 'technique' should be specified as ribose-seq, emRiboseSeq, RHII-HydEn-seq, Alk-HydEn-seq, or Pu-seq. By default, Ribose-Map analyzes all genomic units of the reference genome collectively unless one or more units are specified as 'other' in the configuration file.

**Table 3.** Troubleshooting Advice

| Step(s) | Problem | Possible Reason | Solution |
| --- | --- | --- | --- |
| 1-7 | Modules cannot be located | Path to modules not specified | Specify full path to ribose-map/modules when running any module or directly run module from within ribose-map/modules |
| 1-7 | Software cannot be located | ribosemap_env not activated | Activate software environment:<br>conda activate ribosemap_env |
| 3 | Coordinate Module outputs empty BED file to results folder | Technique parameter is not specified correctly in config file | Technique parameter should be set to ribose-seq, emRiboSeq, RHII-HydEn-seq, Alk-HydEn-seq, or Pu-seq in config file |
| 4-7 | Composition, Sequence, Distribution, and/or Hotspot Modules do not output results | These modules depend on BED file created by Coordinate Module. Thus, BED file must be present in Coordinate Module's results folder for subsequent modules to run | Run Coordinate Module to obtain BED file |

#### Downloading and Preparing Required Data Files

First, the conda software environment should be activated. Next, the FASTQ file of ribose-seq data should be downloaded from NCBI’s Short Read Archive using the fastq-dump tool of the SRA Toolkit. Then, the FASTA and .chrom.sizes files for *S. cerevisiae* should be downloaded from the UCSC genome browser website and the Bowtie 2 indexes should be built from the FASTA file.

Activate conda software environment:

$ conda activate ribosemap_env

Download FASTA file from UCSC genome browser for sacCer2 reference genome:

$ wget http://hgdownload.soe.ucsc.edu/goldenPath/sacCer2/bigZips/sacCer2.fa.gz

Download .chrom.sizes file from UCSC genome browser for sacCer2 reference genome: $ wget http://hgdownload.soe.ucsc.edu/goldenPath/sacCer2/bigZips/sacCer2.chrom.sizes

If the .chrom.sizes file is not readily available for download for the particular genome of interest, see **Box 1** below on creating this file from the FASTA file using SAMtools and Linux commands.

###### Box 1 Create file containing the reference genome chromosome sizes

If the .chrom.sizes file is not readily available for download for the genome of interest, this file can be created from the reference genome FASTA file using the following commands:

$ samtools faidx sacCer2.fa > sacCer2.fa.fai

$ cut -f1,2 sacCer2.fa.fai > sacCer2.chrom.sizes

Unzip FASTA file

$ gunzip sacCer2.fa.gz

Download FASTQ files of rNMP sequencing data from NCBI’s Short Read Archive:

$ fastq-dump SRR11364933

$ fastq-dump SRR1734967

Create Bowtie 2 indexes for reference genome:

$ bowtie2-build sacCer2.fa sacCer2

### PROCEDURE

1. **Trim reads based on quality, length, and adapter content, Timing <5 min** The first 12 nucleotides of the raw ribose-seq reads contain an 11 nucleotide UMI/barcode sequence plus the tagged nucleotide. Therefore, the minimum allowed read length was selected as 62 nucleotides so that at least 50 nucleotides of genomic DNA remain after the UMI/barcode sequence is removed during the Alignment Module. The raw emRiboSeq reads do not contain a UMI or barcode, so the minimum allowed read length was selected as 51 nucleotides (the tagged nucleotide + 50 nucleotides of genomic DNA for alignment). $ trim_galore -a AGTTGCGACACGGATCTATCA -q 15 --length 62 SRR11364933.fq -o output $ trim_galore --illumina -q 15 --length 50 SRR1734967.fq -o output **? TROUBLESHOOTING** **Steps 2-7 should be run from within the ribose-map/modules directory**
2. **Run the Alignment Module, Timing <5 min** $ ribose-map/modules/ribosemap alignment ribose-seq.config $ ribose-map/modules/ribosemap alignment emRiboSeq.config **? TROUBLESHOOTING** See **Box 2** for details about calculating read statistics using files generated from this module.
3. **Run the Coordinate Module, Timing <5 min** $ ribose-map/modules/ribosemap coordinate ribose-seq.config $ ribose-map/modules/ribosemap coordinate emRiboSeq.config **? TROUBLESHOOTING**
4. **Run the Composition Module, Timing <5 min** $ ribose-map/modules/ribosemap composition ribose-seq.config $ ribose-map/modules/ribosemap composition emRiboSeq.config **? TROUBLESHOOTING**
5. **Run the Sequence Module, Timing ~2 hr** $ ribose-map/modules/ribosemap sequence ribose-seq.config $ ribose-map/modules/ribosemap sequence emRiboSeq.config **? TROUBLESHOOTING**
6. **Run the Distribution Module, Timing ~30 min** $ ribose-map/modules/ribosemap distribution ribose-seq.config $ ribose-map/modules/ribosemap distribution emRiboSeq.config **? TROUBLESHOOTING**
7. **Run the Hotspot Module, Timing <5 min** $ ribose-map/modules/ribosemap hotspot ribose-seq.config $ ribose-map/modules/ribosemap hotspot emRiboSeq.config **? TROUBLESHOOTING**

##### Box 2 Calculating read statistics for rNMP sequencing data prepared using barcode and/or UMIs

rNMP sequencing libraries can be prepared using a molecular barcode and/or UMIs. After running Ribose-Map’s Alignment Module, the following commands can be run within the output ‘alignment’ folder within the results folder to determine the percentage of reads that contain the correct barcode and the percentage of reads that remain after de-duplication with UMI-tools.

###### Reads containing barcode (%) =

$ bc -l <<< $(wc -l demultiplexed1.fq | awk ‘{print $1 / 4}’)/$(wc -l SRR11364933_trimmed.fq

| awk ‘{print $1 / 4}’)*100 | xargs printf “%.*f\n” 2

###### Reads after de-duplication (%) =

$ bc -l <<< $(samtools view -c -F 4 SRR11364933.bam)/$(samtools view -c -F 4 sorted.bam)*100

| xargs printf “%.*f\n” 2

**? TROUBLESHOOTING**

Troubleshooting advice can be found in **Table 3**.

**Figure 3.**
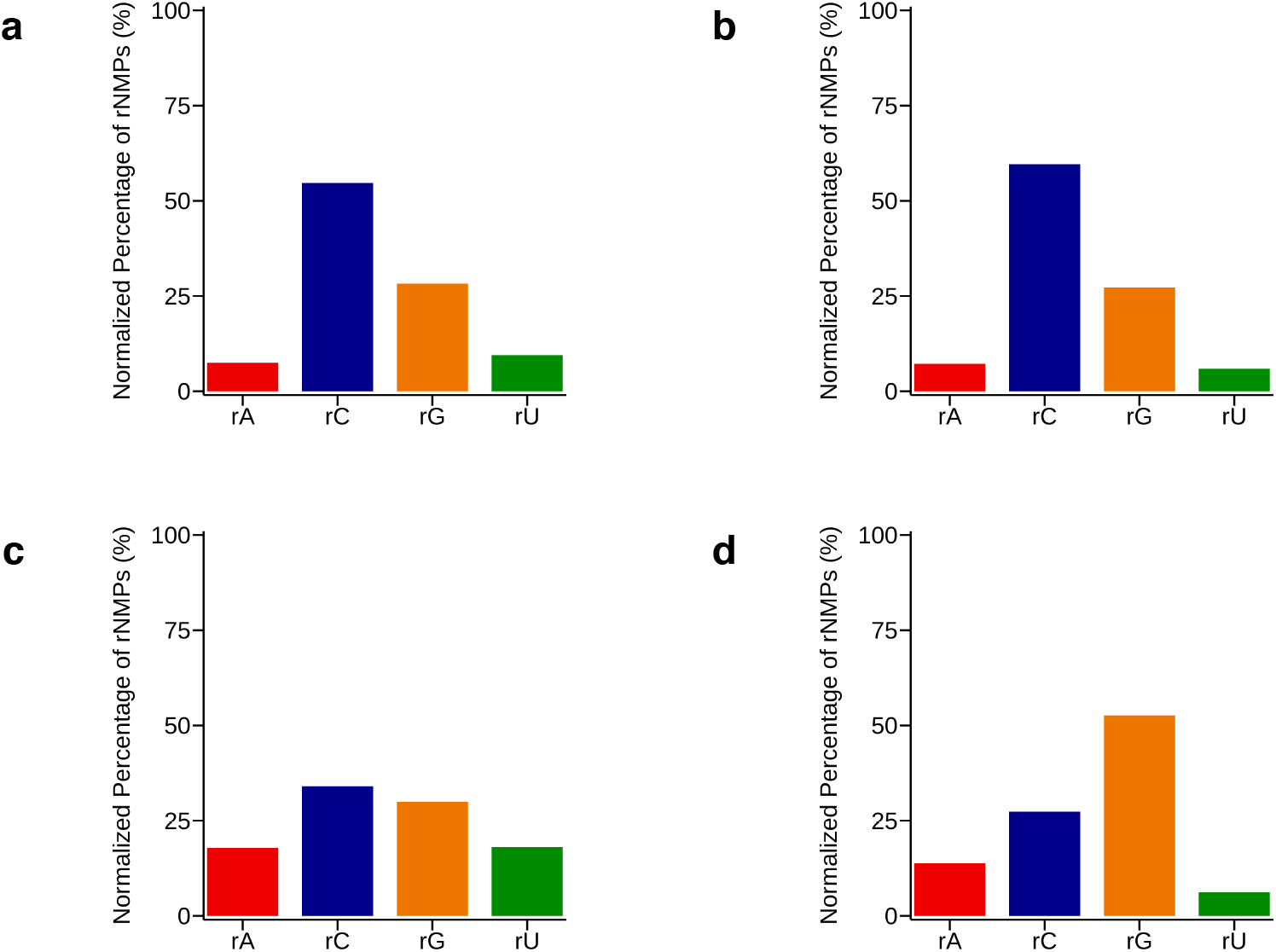
Normalized percentages of rNMP composition in the DNA of RNase H2 defective *S. cerevisiae;* (**a**) nuclear and (**b**) mitochondrial DNA of strain E134 (library SRR11364933) captured using ribose-seq; (**c**) nuclear and (**d**) mitochondrial DNA of strain (Δl−2)l-7BYUNI300 (library SRR1734967) captured using emRiboSeq. Percentages were calculated and plotted using Ribose-Map’s Composition Module and are normalized to the sacCer2 reference genome.

## TIMING

Running this protocol on the *S. cerevisiae* data provided will take ~30 min of hands-on time and 3 hr of computing time for a high-performance computer cluster with 2 nodes, 4 processors per node, and 5 GB of memory per core. For other data sets, the timing may take longer depending on the volume of sequencing data, the size of the reference genome, and the computer used.

## ANTICIPATED RESULTS

The following results were obtained using each module of Ribose-Map for the sample datasets stated previously. Regardless of the dataset, the user should anticipate output files similar to those for the sample datasets used in this protocol.

### Alignment Module

The Alignment Module outputs a Binary Alignment Map (BAM) file of the aligned reads (also de-multiplexed and de-duplicated if needed). Approximately 90% or more of the input reads should align to the reference genome ^13^. Lower alignment rates could indicate low quality reads and/or the presence of contaminants (e.g., custom sequencing adapter) in the reads. In addition, lower alignment rates could result from a substantial level of mismatched nucleotides, insertions, and/or deletions compared to the reference genome, suggesting the reference is not ideal for that particular sample. In addition to the BAM file, the Alignment Module also outputs any relevant intermediate files, including a FASTQ file after the input reads have been de-multiplexed as well as a BAM file after alignment but before de-duplication based on UMIs. Based on these intermediate files, the user can easily track how the Alignment Module processes the input reads and calculate pertinent read statistics, such as the percentage of reads containing the barcode and/or the percentage of reads remaining after PCR de-duplication (See **Box 2**). **Table 4** shows the read alignment statistics for the ribose-seq and emRiboSeq example data sets.

**Table 4.** Read alignment statistics for ribose-seq data (library SRR11364933) and emRiboSeq data (library SRR1734967).

| **Library** | **Aligned Reads (%)** | **Barcoded Reads (%)** | **Unique Reads (%)** |
| --- | --- | --- | --- |
| SRR11364933 | 98.88 | 81.69 | 39.39 |
| SRR1734967 | 98.95 | N/A | N/A |

Aligned Reads represents the Reads aligned to both nuclear and mitochondrial DNA. Barcoded Reads represents the Reads containing barcode specified in configuration file. Unique Reads represents the Reads remaining after reads are PCR de-duplicated based on UMI and genomic coordinates. Barcoded Reads = not applicable (N/A) for emRiboSeq library SRR1734967 since the reads were not barcoded during library preparation. Unique Reads = not applicable (N/A) for emRiboSeq library SRR1734967 since the library was not PCR amplified.

### Coordinate Module

Based on the BAM file output from the Alignment Module, the Coordinate Module outputs a browser extensible data (BED) file of the genomic coordinates of all rNMPs present in the sequencing library (6 columns: chromosome, 0-based start coordinate, 0-based end coordinate, read name, and DNA strand (+/-)). Each line of this file represents the genomic coordinates of an rNMP, and multiple lines may have the same coordinates. The number of aligned reads (concordantly aligned reads for paired-end data) in the BAM file should be equal to the number of rNMPs in the BED file unless some reads were removed during quality control for emRiboSeq, Alk-HydEn-seq, or Pu-seq data as described previously (see Locating rNMPs with the Coordinate Module section of Experimental Design) or if some of the reads did not meet the quality threshold specified in the configuration file. In addition, the Coordinate Module also outputs a tab-delimited file of unqiue rNMP sites and counts of rNMPs at each site as well as BED files of rNMP genomic coordinates for each genomic unit specified in the configuration file. **Table 5** shows the total number of rNMPs and the number of unique rNMP sites in the mitochondrial DNA of the ribose-seq and emRiboSeq example data sets.

**Table 5.** rNMP genomic coordinate statistics for mitochondrial and nuclear ribose-seq data (library SRR11364933) and emRiboSeq data (library SRR1734967).

| Library | Nucleus |  | Mitochondria |  |
| --- | --- | --- | --- | --- |
|  | Total rNMPs | rNMP Sites | Total rNMPs | rNMP Sites |
| SRR11364933 | 693,359 | 388,437 | 48,131 | 13,335 |
| SRR1734967 | 2,745,709 | 2,280,795 | 47,037 | 25,102 |

Total rNMPs represents the Number of rNMPs in library after alignment, de-multiplexing, and de-duplication (if required). rNMP Sites represents the Number of genomic coordinates where at least one rNMP is present.

### Composition Module

One of the first steps that should be taken to identify possible biological signatures of rNMP incorporation is determining if there are any biases in the composition of rNMPs incorporated into the genome of interest, meaning the relative frequencies of the four types of rNMPs (r[A, C, G, U]MP) that can be incorporated into the genome. The Composition Module outputs the raw counts of each type of rNMP and the percentages of each rNMP normalized to the corresponding nucleotide frequencies of the reference genome. The counts and frequencies are calculated for each genomic unit specified in the configuration file. In addition, the Composition Module plots the normalized percentages of rNMP composition. Based on these plots, the user can readily determine if certain types of rNMPs are preferentially incorporated into the genome of interest. In this example, rCMP is the most frequently incorporated rNMP relative to the reference genome in the mitochondrial DNA of the ribose-seq data set and rGMP is the most frequently incorporated rNMP in the mitochondrial DNA of the emRiboSeq data set, while rUMP and rAMP are the least frequently incorporated rNMPs in both of these data sets (**Figure 3**).

### Sequence Module

To reveal if rNMP incorporation is influenced by DNA sequence context, the Sequence Module calculates and plots the frequencies of each type of nucleotide (A, C, G, and U/T) at sites of rNMP incorporation and surrounding those sites. The Sequence Module calculates and plots these frequencies for all types of rNMPs combined as well as for each rNMP individually. In this example, rNMPs are more frequently incorporated into GC-rich regions of *S. cerevisiae* mitochondrial DNA than would be expected given the nucleotide frequencies of the reference genome for all four types of rNMPs r[A, C, G, U]MP combined (**Figure 4**). By examining the DNA sequence context for each type of rNMP individually, the user can explore if the GC-rich sequence context is specific to all types of rNMPs or a subset. Based on these individual plots, it becomes clear that although all four types of rNMPs are more frequently incorporated up/downstream from deoxyribonucleotide C and deoxyribonucleotide G, rCMP and rGMP are incorporated into GC-rich areas of the mitochondrial DNA at a much higher frequency than rAMP and rUMP.

**Figure 4.**
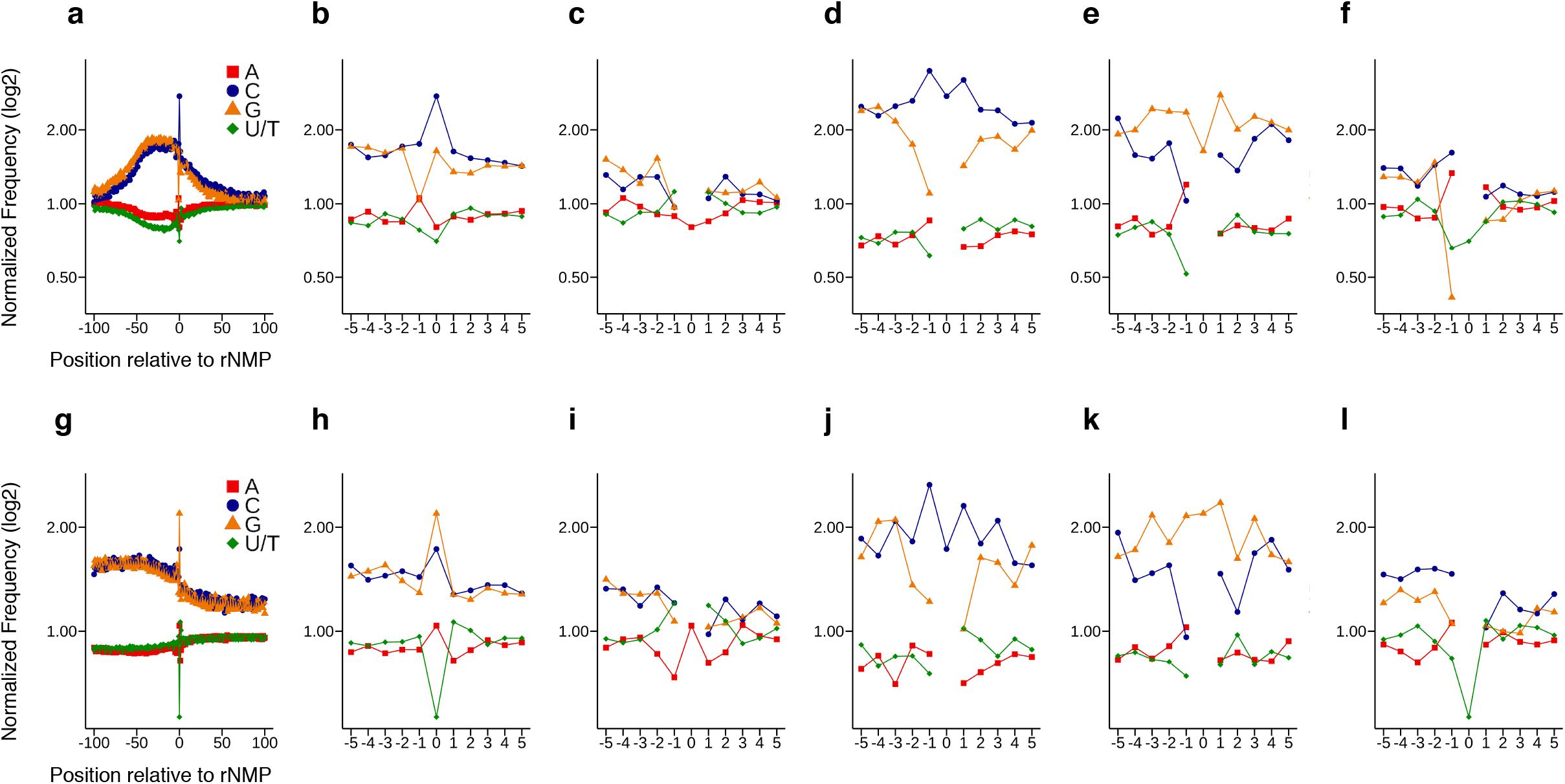
DNA sequence context of rNMPs in the mitochondrial DNA of *S. cerevisiae* (**a-f**) strain E134 (library SRR11364933) captured using ribose-seq and (**g-l**) strain (Δl−2)l-7BYUNI300 (library SRR1734967) captured using emRiboSeq and obtained using Ribose-Map’s Sequence Module. Ribose-seq data (**a**) zoomed-out for r[A, C, G, U]MP combined, (**b**) zoomed-in for r[A, C, G, U]MP combined, (**c**) zoomed-in for rAMP, (**d**) zoomed-in for rCMP, (**e**) zoomed-in for rGMP, and (**f**) zoomed-in for rUMP. emRiboSeq data (**g**) zoomed-out for r[A, C, G, U]MP combined, (**h**) zoomed-in for r[A, C, G, U]MP combined, (**i**) zoomed-in for rAMP, (**j**) zoomed-in for rCMP, (**k**) zoomed-in for rGMP, and (**l**) zoomed-in for rUMP. Nucleotide frequencies are normalized to those of the sacCer2 reference genome and are shown relative to rNMP sites (position 0 on x-axis = rNMP).

### Distribution Module

To assess the genome-wide distribution of rNMP incorporation, the Distribution Module outputs tab-delimited files containing the per-nucleotide coverage of rNMPs across each genomic unit of interest (e.g., chromosomes, mitochondria, plasmids) normalized to reads per million (RPM) to allow data sets of different sequencing depth to be readily compared (**Figure 5**). In addition, the Distribution Module outputs BedGraph files of rNMP coverage that can be uploaded to any genome browser, such as the UCSC genome browser, for further investigation.

**Figure 5.**
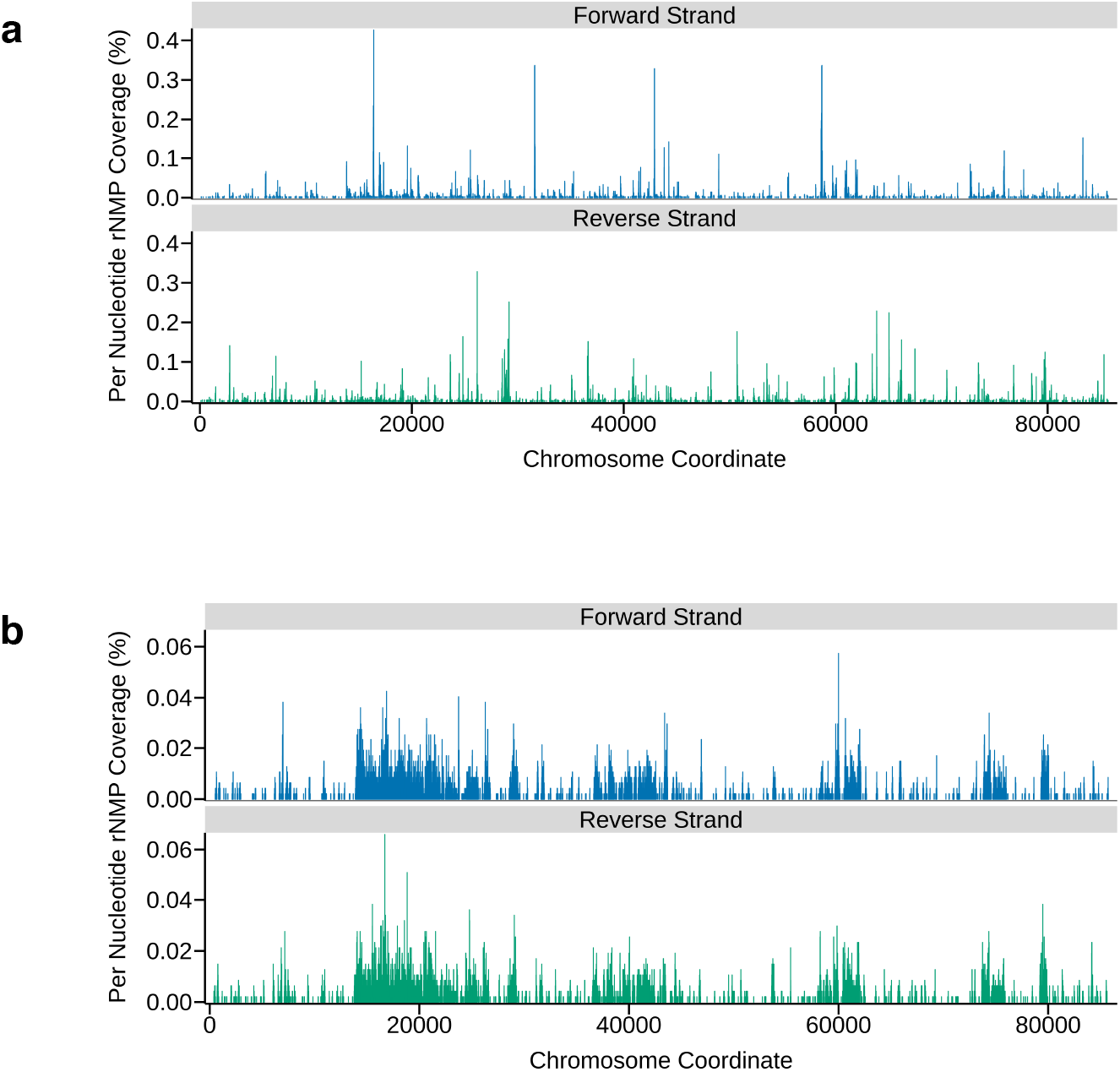
Distribution of rNMPs in the mitochondrial DNA of *S. cerevisiae* (**a**) strain E134 (library SRR11364933) captured using ribose-seq and (**b**) strain (Δl−2)l-7BYUNI300 (library SRR1734967) captured using emRiboSeq and obtained using Ribose-Map’s Distribution Module. Per-nucleotide coverage of rNMPs is normalized to reads per million (RPM).

### Hotspot Module

To identify sites that are susceptible to rNMP incorporation and determine if these sites share a consensus sequence, the Hotspot Module outputs the genomic coordinates of the top 1% most abundant sites of rNMP incorporation and plots their consensus sequence (**Figure 6**). These hotspots can also be cross-referenced with gene annotations to explore genetic or epigenetic consequences of rNMP incorporation in the genomic DNA of interest.

**Figure 6.**
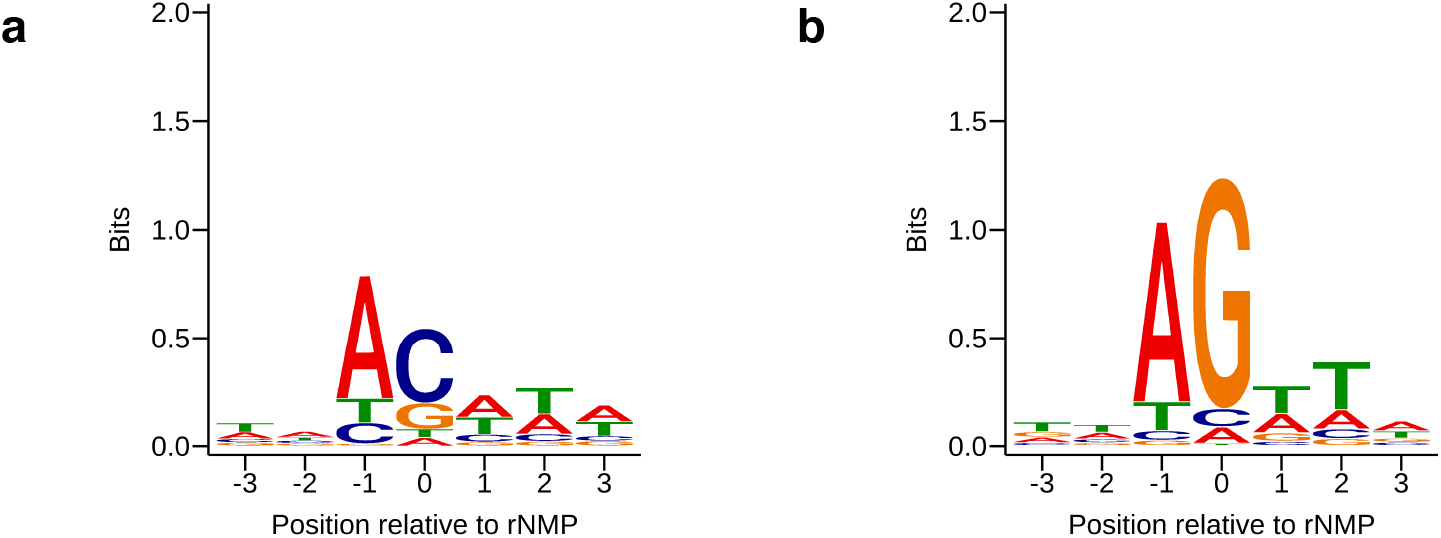
Consensus sequences of the top 1% most abundant sites of rNMP incorporation in the mitochondrial DNA of *S. cerevisiae* (**a**) strain E134 (library SRR11364933) captured using ribose-seq and (**b**) strain (Δl−2)l-7BYUNI300 (library SRR1734967) captured using emRiboSeq and obtained using Ribose-Map’s Hotspot Module (position 0 on x-axis = rNMP).

## Author Contributions

A.L.G. is the lead developer of Ribose-Map and wrote the manuscript with input from F.S. A.L.G. and along with F.S. designed and developed the protocol and generated the example experiments.

## ACKNOWLEDGMENTS

We are grateful to P. Xu, T. Yang, K. Mukherjee, and D. Kundnani for suggestions on this manuscript as well as M. Borodovsky, I.K. Jordan, S. Yi, F. Vannberg and all members of the Storici laboratory for their advice during the course of this study. This research was supported by the National Institutes of Health [R01ES026243-01 to F.S.]; the Parker H. Petit Institute for Bioengineering and Bioscience at Georgia Institute of Technology [#12456H2 to F.S.], and the Howard Hughes Medical Institute Faculty Scholar grant [#55108574 to F.S.].

## Competing financial interests

The authors declare that they have no competing financial interests.

## Notes

### Competing Interest Statement

The authors have declared no competing interest.

